# Preclinical model for the study of immune responses specific for a hepatic self-antigen

**DOI:** 10.1101/2022.06.22.497185

**Authors:** Anaïs Cardon, Jean-Paul Judor, Thomas Guinebretière, Richard Danger, Arnaud Nicot, Sophie Brouard, Sophie Conchon, Amédée Renand

**Author notes:** Equal contribution (Sophie Conchon, Amédée Renand). Corresponding author: Dr Amédée Renand, INSERM UMR1064 – Center for Research in Transplantation and Translational Immunology, CHU Nantes – Hôtel Dieu, 30 boulevard Jean Monnet, 44093, Nantes Cedex 01, France.

## Abstract

The liver displays a strong capacity to induce tolerance toward hepatic antigens. However, hepatic tolerance can be overcome with the development of local autoimmune diseases such as autoimmune hepatitis (AIH). This chronic inflammatory disorder leads to a progressive destruction of liver parenchyma if non-treated. Although the CD4^+^ T cell response seems a key player of this immune disorder, the dynamics and biology of emerging liver antigen-specific CD4^+^ T cells are poorly described. Here, we developed a new murine model which mimics hepatic autoreactivity allowing the study and monitoring of antigen-specific CD4^+^ T cells from their emergence to local immune response. We show that the induction of the expression of an antigen in the liver in non-inflammatory condition leads to antigen tolerance. In inflammatory condition, using viral vector transduction, we observe the development of a complete adaptive immune response concomitant with the antigen expression in the liver. The presence of antigen-specific CD4^+^ T cells in the liver is associated to transient hepatic damages. Interestingly, the neo-antigen expression by hepatocytes after peripheral immunisation induces the recruitment of antigen-specific CD4^+^ T cells and hepatic damages. These data demonstrate that the recruitment of antigen-specific CD4^+^ T cells in the liver is conditioned by an immune coordination between surface antigen expression by hepatocytes and peripheral immune response and mimics the first step of a local autoreactive process. In the long-term, we observe that the hepatic environment has the capacity to control the local, but not the systemic, antigen-specific CD4^+^ T cells. Additional immune events might be involved in the long-term chronic immune reactivity in the liver, following the first steps described in this study.

**Key points:**

- Antigen expression in the liver in non-inflammatory condition leads to antigen tolerance
- Antigen expression in the liver in immunization condition (concomitant or pre-existing) is sufficient to induce liver recruitment of antigen-specific CD4^+^ T cells and hepatic damages
- This model mimics the first step of an autoreactive process against a liver antigen
- In the long-term, the hepatic environment induces a local tolerance toward the antigen expression and the clearance of liver-infiltrating, but not peripheral, antigen-specific CD4^+^ T cells
- This new murine model can be of interest for the analysis of the immunomodulatory pathways implicated in liver tolerization of autoreactive CD4^+^ T cells and identification of potential extrinsic factors implicated in an acute-to-chronic transition as observed during autoimmune hepatitis

## Introduction

The liver displays strong tolerogenic properties mediated by both parenchymal hepatocytes and non-parenchymal cells (NPCs). These cells, particularly liver sinusoidal endothelial cells, Kupffer cells and hepatocytes, are able to act as antigen presenting cells to promote activation of T cells^1^. In normal condition, this activation leads to an anergic phenotype^2,3^, differentiation into regulatory T cells (Tregs)^4,5^ or even apoptosis^6,7^.

However, the robust liver capacity to induce tolerance can be overcome leading to the development of autoimmunity. Autoimmune hepatitis (AIH) is a chronic inflammatory disorder causing a progressive destruction of liver parenchyma^8^. It is a rare worldwide disease with a prevalence of 16.9 per 100,000 in Europe and North America and an incidence of 0.1 to 1.9 per 100,000 per year^9^. This pathology is characterized by the presence of autoantibodies, elevated immunoglobulin G (IgG) level in the sera of patients, and a typical histological feature in the liver, the interface hepatitis. This important lymphocytic infiltration in the liver is composed mainly of CD4^+^ T cells but also of CD8^+^ T cells and B cells^8^. AIH is divided into two types, depending on autoantibodies present in patients’ blood: type 1 AIH (anti-nuclear antibodies, anti-smooth muscle antibodies and anti-soluble liver antigen – SLA – antibodies) and type 2 AIH (anti-liver kidney microsomal type 1 – LKM-1 – antibodies and anti-liver cytosol – LC-1 – antibodies). AIH is a multifactorial disease associating genetic predispositions, affecting genes coding for class II molecules of the major histocompatibility complex (MHC-II), and environmental factors, such as drug exposure and molecular mimicry^8^. Patients are treated with immunosuppressive drugs (prednisolone associated or not with azathioprine) but struggle to reach long-term remission^10,11^.

Preponderance of CD4^+^ T cells in the hepatic infiltrate, genetic predisposition affecting genes coding for MHC-II molecules, involved in antigen presentation to CD4^+^ T cells, and accumulation of autoantibodies, generated after interaction between B cells and CD4^+^ T cells, point toward a major role of autoreactive CD4^+^ T cells in the immunopathogenesis of AIH. Data obtained in AIH patients show an enrichment of pro-inflammatory CD4^+^ T cells in liver and blood and defective regulatory mechanisms^8,12^. Particularly, frequencies of pro-inflammatory Th1 and Th17 cells and their related cytokines (IFN-γ, TNF-α, IL-17) are increased in patients ^13–15^. Whether patients present quantitative and/or qualitative CD4^+^ Tregs defects is still debated^15–18^. Recently, our team observed that, despite immunosuppressive treatment, AIH patients display a persistent immune dysregulation in their blood with a residual CD4^+^ T cell infiltrate in the liver^12^. By tracking SepSecs-specific CD4^+^ T cells in patients with anti-SLA antibodies, we identified and provided a deep characterization of a unique autoreactive CD4^+^ T cell population with a pro-inflammatory/B-helper profile (PD-1^+^ CXCR5^-^ CCR6^-^ CD27^+^ IL-21^+^). This subset is enriched in the blood of AIH patients independently of the presence of the anti-SLA antibodies^19^, and it represents the major reservoir of autoreactive CD4^+^ T cells in patients. Therefore, the tracking of these cells associated to the analysis of the mechanisms involved in their emergence could provide new therapeutical target for AIH treatment.

As diagnosis generally occurs quite late after disease onset, and because AIH is a chronic disease with long-term evolution, the study of the early events implicated in the autoreactive CD4^+^ T cells generation cannot be performed in patients, thus murine models represent valuable tools for this purpose. Murine models of AIH, based on immunisation using viral vectors encoding for human CYP2D6 or FTCD proteins^20,21^, have underlined the importance of molecular mimicry and genetic susceptibility for the development of murine AIH. However, emergence and biology of antigen-specific autoreactive CD4^+^ T cells involved in the first steps of the adaptive immune response against an autoantigen expressed in the liver have not been studied. In this study, we propose a new murine model which mimics hepatic autoreactivity. In this model, we can induce hepatic expression of a model antigen, hemagglutinin (HA), in different conditions and follow the emergence of antigen-specific responses, especially antigen-specific CD4^+^ T cells.

## Methods

### Viral vectors

Adenoviral CAG Cre vectors (Ad Cre) were produced with a vector plasmid containing the Cre recombinase coding sequence inserted behind the CAG promoter and followed by a poly-A signal. Control adenoviral CMV GFP vector (Ad Ct) consisted of the GFP coding sequence inserted between the CMV promoter and a poly-A signal.

Ad Cre and Ad Ct productions were performed by the INSERM UMR 1089 Centre de Production de Vecteurs (Nantes, France).

### Mice

Heterozygous TTR-Cre inducible mice (Cre^ind-/+^)^22^ were back-crossed on a Balb/c background for at least 10 generations (TAAM, CDTA CNRS Orléans). They were cross-bred with homozygous Rosa26 HA floxed mice (HA^fl^)^23^ resulting in HA^fl^/Cre^ind-^ mice and HA^fl^/Cre^ind+^ mice. Male and female eight to twelve-week-old mice were used for each experiment. All mice were housed at the UTE IRS-UN animal facilities (Nantes, France) where they were fed *ad libitum* and allowed continuous access to tap water. Procedures were approved by the regional ethical committee for animal care and use and by the Ministère de l’enseignement supérieur et de la recherche (agreements APAFIS #2054 and #28582). All experiments were performed in accordance with relevant guidelines and regulations.

For the induction of HA expression in hepatocytes in non-inflammatory condition, normal diet of HA^fl^/Cre^ind-^ mice (control) and HA^fl^/Cre^ind+^ mice were substituted by tamoxifen dry food (0.5g/kg tamoxifen + 5% saccharose; Safe, France) for 14 days in free access.

For the induction of HA expression in hepatocytes in inflammatory condition, HA^fl^/^ind-^ mice and HA^fl^/Cre^ind+^ mice were injected intravenously with 3×10^9^ ip (infectious particle) per mice of the Ad Cre vector or of the Ad Ct vector for the control group.

For the peripheral immunization against HA, mice HA^fl^/Cre^ind-^ mice and HA^fl^/Cre^ind+^ mice were injected intramuscularly with 1.5×10^9^ ip per mice of the Ad Cre vector or of the Ad Ct vector for the control group.

### Genotyping

Small tail biopsies were taken from 3 week-old mice to perform Cre genotyping. Briefly, samples were digested overnight at 56°C in 100μL of TNT-PK buffer (TNT: Tris HCl pH 8.5 50mM; NaCl 100mM;Tween 20 0.5% *I* proteinase K 0.2mg/mL). PK were inactivated at 95°C during 15min. 60ng of DNA was used for PCR mix reaction prepared according to the manufacturer protocol (Herculase II Fusion DNA Polymerase, Agilent). The following Cre primers were used to carry out the PCR amplification: forward primer 5’-CCTGGAAAATGCTTCTGTCCG-3’, reverse primer 5’-CAGGGTGTTATAAGCAATCCC-3’. Amplification program was run on a Veriti Thermal Cycler (Applied Biosystems; Foster City, USA) and consist of 1 cycle at 95°C for 2min, 35 cycles of 95°C for 20sec, 60°C for 20sec and 72°C for 30sec, followed by 1 cycle at 72°C for 3min. Finally, PCR products are visualized on Caliper LabChip (PerkinElmer).

### RNA extraction, reverse transcription and quantitative PCR

Total RNA was extracted from organ tissue using TRIzol^TM^ reagent (ThermoFisher Scientific #15596026) and purified with the QiagenRNeasy Mini Kit (Qiagen #74106) according to the manufacturer’s protocol.

Reverse transcription was performed using 2μg of total RNA mixing with poly-dT24 20μg/mL (Eurofins Genomics), DTT 8mM (ThermoFisher Scientific #18057018) and dNTP 20mM (ThermoFisher Scientific #10297018) and incubated at 70°C during 10min followed by 4°C for 5min. Then, first strand buffer 1X (ThermoFisher Scientific #18057018), M-MLV reverse transcriptase 200U (ThermoFisher Scientific #18057018) and RNAse OUT inhibitor 40U (ThermoFisher Scientific #10777019) were added and incubated at 37°C for 1h, followed by 15min at 70°C.

Real-time RT-PCR was performed using the ViiA^TM^ 7 Real-Time PCR System and Power SYBR^TM^ Green PCR Master Mix (ThermoFisher Scientific #4368708). Primers for HA (forward 5’-AAACTCTTCGCGGTCTTTCCA-3’; reverse 5’-GATAAGGTAGCTTGGGCTGC-3’) and β-actine (forward 5’-TACCACAGGCATTGTGATGG-3’; reverse 5’-AATAGTGATGACCTGGCCGT-3’) were used for detection. Relative gene expressions were calculated by the 2^-ΔΔCt^ method.

### Western blot

Total proteins were extracted from liver samples via RIPA buffer treatment. 25μg of protein were denatured at 95°C for 5min in Laemmli Sample Buffer (BIORAD #1610747) with DTT 0.1M. Preparation was separated by SDS-PAGE on Mini-PROTEAN TGX Precast Protein Gels (BIORAD #4561036) in migration buffer (Tris base 15g/l, glycine 72g/l, SDS 10g/l) and transferred onto a PVDF membrane with the Trans-Blot Turbo Transfer System (BIORAD). The membrane was blocked using a blocking solution (TBS; Tween 20 0.1%; skim milk 5%) for 2h then incubated with primary antibodies (1:5000) overnight: anti-HA antibody (rabbit polyclonal antibody; Sinobiological #11684-T62) and anti-β-actin antibody (mouse monoclonal antibody; Cell Signaling #3700). The membrane was washed with TBS – Tween 20 0.1%. HRP-conjugated donkey anti-rabbit IgG (H+L) (1:5000; Cliniscience #E-AB-1080-120) and HRP-conjugated goat anti-mouse IgG + IgM (H+L) (1:10000; Jackson ImmunoResearch #115-036-068) were used to detect rabbit and mouse antibodies respectively during 1h. Secondary antibodies were detected using electrochimioluminescence super signal West Pico (ThermoFisher Scientific #34577) according to the manufacturer instructions. Imaging and analysis of western blots were performed on the ChemiDoc MP Imaging System (BIORAD).

### Cell preparation

Splenocytes were isolated by a mechanical dissociation of spleen in red blood cell lysis buffer (NH_4_Cl 155mM; KHCO_3_ 10mM; EDTA 1mM; Tris 17mM per 1L of sterilized water) before centrifugation.

Liver non-parenchymal cells (NPCs) were isolated as previously described^24^. Briefly, after perfusion with HBSS 1X buffer (ThermoFisher Scientific #14175129), livers were digested with collagenase IV (Sigma-Aldrich #C5138) and NPCs enriched by Percoll (Sigma-Aldrich #GE17-0891-01) density gradient centrifugation and red blood cells lysis.

### Tetramer enrichment and staining

20.10^6^ splenocytes and 3.10^6^ NPCs were stained with both I-A^d^-HA peptide (HNTNGVTAACSHE) and I-E^d^-HA peptide (SFERFEIFPKE) PE-labelled tetramers (NIH Tetramer Core Facility; Atlanta, USA) at room temperature during 1h. Then, cells were washed using PBE buffer (PBS 1X; BSA 0.1%; EDTA 2mM) and stained with magnetic anti-PE microbeads (Miltenyi #130-048-801) at 4°C during 15min. Cells were washed and enriched using magnetic MS columns (Miltenyi #130-042-201). A viability staining was then performed on the positive fraction containing tetramer-enriched cells by incubating cells with 100μL of LIVE/DEAD^TM^ Fixable Aqua Dead Cell Stain kit (ThermoFisher Scientific #L34957) for 15min at 4°C, protected from light. Cells were washed and an extracellular staining were performed using 100μL of fluorescent antibodies for 20min at 4°C, protected from light. For FoxP3 intracellular staining, cells were next permeabilized and fixed for 30min according to the eBioscience^TM^ Foxp3/Transcription Factor Staining Buffer Set (ThermoFisher Scientific #00-5523-00). Fluorescent antibodies targeting the FoxP3 marker were diluted in the permeabilization buffer and incubated with the fixed cells for 1h, protected from light.

The following antibodies were used to perform an extracellular staining: CD4/PerCP-Cy5.5 (clone RM4-5; BD #550954), CD44/APC (clone IM7; BD #559250), PD-1/BV421 (clone J43; BD #562584), CD25/PE-Cy7 (clone PC61; BD #552880) and CD19/BV510 (clone ID3; BD #562956). For the intracellular staining, the FoxP3/AF488 (clone FJK16S, eBioscience #53-5773-82) antibody was used. HA-specific CD4^+^ T cells were defined as: LIVE/DEAD^-^ CD19^-^ CD4^+^ CD44^high^ tetramers^+^ cells.

For cell phenotyping, fluorescence was measured with a BD FACS Canto II (BD Biosciences; Mountain View, USA). FlowJo software was used to analyze data after eliminating doublets and viability^+^ dead cells.

### ELISA test

For the detection of anti-HA antibodies, wells were coated with HA protein 1μg/mL (Sino Biological #11684-V08H) diluted in coating buffer (Na_2_CO_3_ 0.05M; NaHCO_3_ 0.05M; pH 9.2) and incubated at 4°C overnight. Wells were then washed with washing buffer (PBS 1X; Tween 20 0.05%; in distilled water) and saturated with dilution buffer (PBS 1X; Tween 20 0.05%; BSA 1%; in distilled water) for 2h at 37°C. After washing, diluted sera samples (1:500) were added and incubated at 37°C during 2h. Wells were washed and diluted peroxidase goat anti-mouse IgG+IgM (H+L) detection antibody (1:2000; Jackson ImmunoResearch #115-036-068) was added before incubating at 37°C for 1h. Finally, wells were incubated with TMB Substrate Reagent (BD #555214) and reaction was stopped with H_2_SO_4_ 0.5M. The absorbance at 450nm was determined using a Spark^®^ 10M Infinite M200 Pro plate reader (TECAN).

### Transaminase dosage and histology

Dosage of plasma levels of aspartate aminotransferase (AST) and alanine aminotransferase (ALT) was performed by the Centre Hospitalier Universitaire of Nantes (France).

Histological analyses were performed on paraformaldehyde (PFA)-fixed/paraffin-embedded liver samples. Liver lobes were fixed in PFA 4% for 24h at room temperature. Samples were then dehydrated in absolute ethanol then cleared in isopropanol and finally included paraffin. Paraffin-embedded sections (3μm) were stained with hematoxylin-phloxine-saffron (HPS). Paraffin-embedding and HPS staining were performed by the IBISA MicroPICell facility (Biogenouest; Nantes, France). Slides were observed using NanoZoomer (HAMAMATSU) and NDP Scan software.

### Statistical analysis

Prism (GraphPad version 6.01 Software, Inc.) was used for statistical analyses. Group comparison analyses were assessed using non-parametric Kruskal-Wallis tests or one-way ANOVA statistical test. p values < 0.05 were considered statistically significant between two groups (* = p <0.05, ** = p <0.01, *** = p <0.001, **** = p < 0.0001).

## Results

### Concomitant inflammation and antigen expression in the liver induce a local recruitment of antigen-specific CD4^+^ T cells with increased hepatic damages

In order to analyse the dynamics of the emergence of liver antigen-specific CD4^+^ T cells, we used a liver-restricted HA expression model and tracked HA-specific CD4^+^ T cells with specific MHC class II tetramers.

We developed two strategies of induction of the expression of the HA antigen in the liver, based on the Cre/LoxP system. Homozygous Rosa26 HA floxed (HA^fl^) mice were crossed with heterozygous TTR-Cre inducible (Cre^ind^) mice, expressing an inducible Cre recombinase under the control of the hepatocyte-specific promotor (transthyretin, TTR). Both strains were on Balb/c (H-2^d^) background. The feeding of the HA^fl^/Cre^ind+^ offspring with tamoxifen-mixed dry food leads to the expression of HA at the surface of the hepatocytes under non-inflammatory condition. The second strategy consists in the intravenous (i.v.) injection to all offspring (HA^fl^/Cre^ind-^ and HA^fl^/Cre^ind+^) of an adenoviral vector encoding for the Cre recombinase (Ad Cre), which transduces preferentially hepatocytes due to its strong hepatic tropism. This results in the expression of HA at the surface of hepatocytes under inflammatory condition, bypassing the endogenous expression of the inducible Cre in mice. As control, a peripheral model of immunization by intramuscular (i.m.) injection of Ad Cre which only induces local HA expression in the muscle was used (Figure 1A).

**Figure 1:**
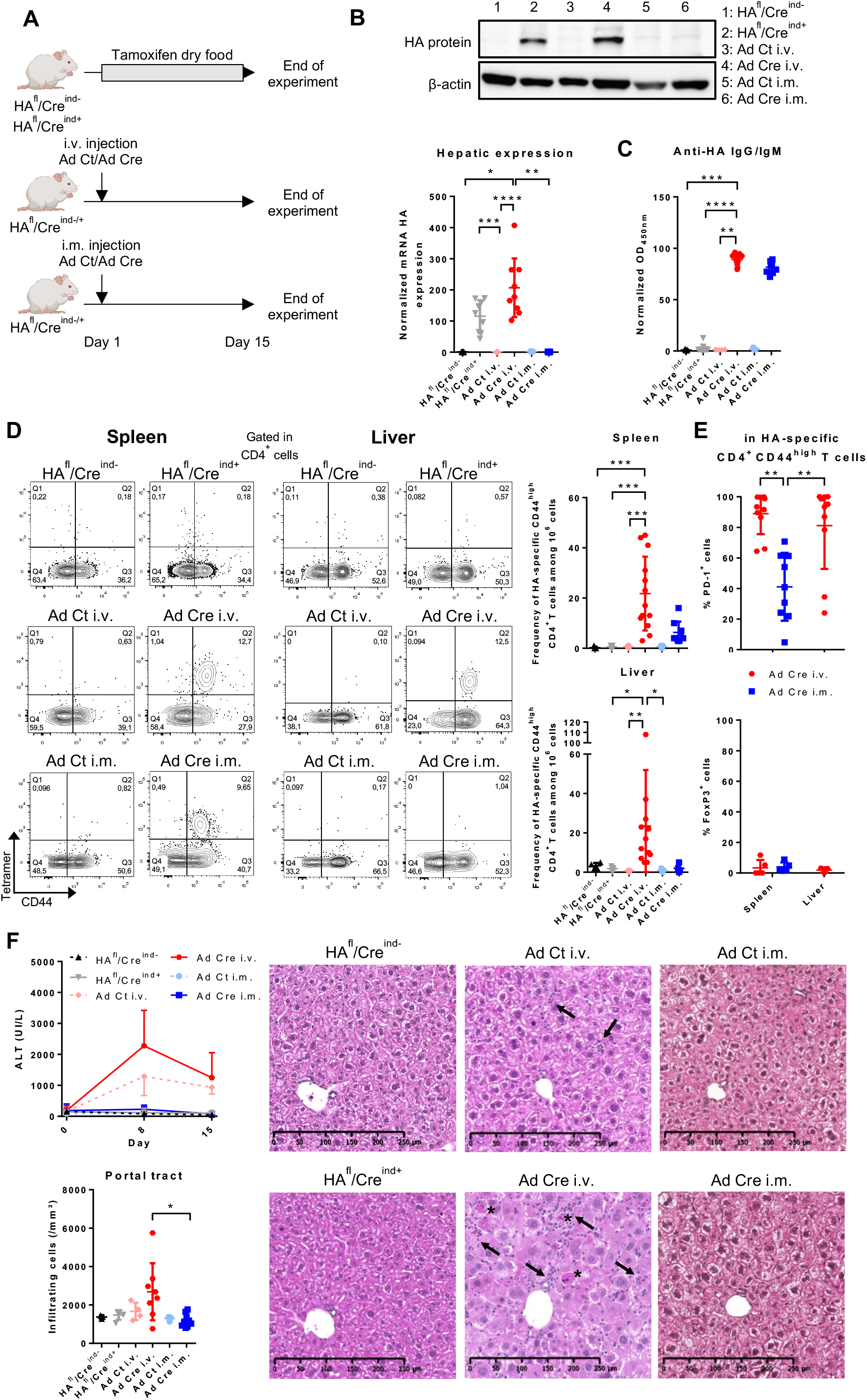
Antigen-specific CD4^+^ T cell response is modulated not only by the localisation of antigen expression but also by the environment. (A) HA^fl^/Cre^ind-^ mice (n = 5) and HA^fl^/Cre^ind+^ mice (n = 9) are fed with tamoxifen dry food (0,5g/kg) for 14 days. HA^fl^/Cre^ind-/+^ mice receive a single i.v. injection (3.10^9^ ip) of Ad Ct (n = 7) or Ad Cre (n = 13). HA^fl^/Cre^ind-/+^ mice receive a single i.m. injection (1,5.10^9^ ip) of Ad Ct (n = 3) or Ad Cre (n = 9). Mice are sacrificed at day 15. (B) Quantitative RT-PCR analysis of HA mRNA expression in liver samples. Total protein from liver samples were incubated on western blot membranes with anti-HA antibodies to detect HA protein expression. β-actine was used as loading control. (C) At the end of experiment, sera were diluted and assayed for anti-HA IgG and IgM. (D) Total splenocytes and liver NPCs are stained with a MHC class II tetramer loaded with HA peptide before a tetramer enrichment step. Cells are stained (Live/Dead Aqua, CD4, CD44, PD-1, CD25, FoxP3, CD19) and analysed by flow cytometry. Representative MHC class II/HA peptide tetramer and CD44 co-staining, gated in live CD19^-^ CD4^+^ cells. Frequency of HA-specific CD4^+^ CD44^high^ T cells are calculated among 10^6^ total cells. (E) Percentage of PD-1^+^ and FoxP3^+^ cells in HA-specific CD4^+^ CD44^high^ T cells are represented. (F) Plasma samples were collected before the start of the experiment and at day 8 and day 15 for dosage of ALT. At the end of the experiment, paraffin-embedded liver sections are stained with HPS coloration to analyse liver infiltration (count of infiltrating cells around portal tracts) and morphology (x10; arrows point lymphocytic infiltration; stars point necroinflammatory activities). All results are normalized and represent the mean (+ SD). p values were calculated using non-parametric Kruskal-Wallis test, * = p <0.05, ** = p <0.01, *** = p <0.001, **** = p < 0.0001.

After two weeks, the HA expression restricted to the liver can be detected both in tamoxifen-fed HA^fl^/Cre^ind+^ mice and in Ad Cre i.v.-injected HA^fl^/Cre^ind-^ and HA^fl^/Cre^ind+^ mice (termed as HA^fl^/Cre^ind-/+^ mice) (Figure 1B, Supplementary figure 1, Supplementary figure 2). Expression of HA in Ad Cre i.m. mice was restricted to the muscle. In control tamoxifen-fed HA^fl^/Cre^ind-^ mice and Ad Ct i.v. or i.m. groups, no expression of HA was observed in the liver and in any other organs (Figure 1B, Supplementary figure 1, Supplementary figure 2).

As expected, when mice are immunized by i.m. injection of Ad Cre, the induction of HA expression in the muscle in this inflammatory condition leads to the generation of a HA-specific humoral response, as demonstrated by the presence of anti-HA IgG and IgM (81.65 ± 5.72 arbitrary units, a.u.) (Figure 1C, Supplementary figure 2). Interestingly, in Ad Cre i.v. mice, the hepatic expression of HA under inflammatory condition leads to a comparable HA-specific humoral response (89.00 ± 4.95 a.u.). On the contrary, tamoxifen fed HA^fl^/Cre^ind+^ mice did not develop humoral HA-specific response, despite hepatic expression of HA. No anti-HA specific antibodies were detected in control mice (tamoxifen-fed HA^fl^/Cre^ind-^ mice and Ad Ct i.v. or i.m. mice) (Figure 1C, Supplementary figure 2).

The HA-specific CD4^+^ T cell response was tracked with MHC class II tetramer loaded with HA peptides. HA-specific CD4^+^ T cells were detected neither in the spleen nor in the liver of tamoxifen-fed HA^fl^/Cre^ind+^ mice (Figure 1D). However, HA-specific CD4^+^ CD44^high^ T cells were detected in the spleen of both Ad Cre i.m. mice (frequency per 10^6^ cells: 6.33 ± 4.27) and Ad Cre i.v. mice (frequency per 10^6^ cells: 21.77 ± 14.68), while none were detected in their respective control (Ad Ct) (Figure 1D, Supplementary figure 2). Liver HA-specific CD4^+^ CD44^high^ T cells were detected only in Ad Cre i.v. mice (frequency per 10^6^ cells: 23.50 ± 28.38) (Figure 1D, Supplementary figure 2). This data demonstrates that our model allows the generation of a complete adaptive immune response against a liver antigen under inflammatory condition with local recruitment of antigen-specific CD4^+^ T cells.

However, the liver has strong tolerogenic properties, which have been shown to involve the induction of regulatory T cells. To understand if our model induces the generation/expansion of regulatory CD4 T cells or of pro-inflammatory CD4 T cells, we performed a comparison of the immune phenotype of the HA-specific CD4^+^ CD44^high^ T cells isolated from Ad Cre i.v. vs Ad Cre i.m. mice. This analysis revealed that these HA-specific CD4^+^ T cells are CD25^-^ FoxP3_-_ in both groups, but that those from Ad Cre i.v. mice contain a higher proportion of PD-1^+^ (CD279) cells (spleen: 88.98 ± 13.41%; liver: 81.11 ± 28.29%) than those isolated from Ad Cre i.m. mice (spleen: 41.12 ± 22.21%) (Figure 1E, Supplementary figure 3). PD-1^+^ cells were enriched in HA-specific CD4^+^ T cells compared to total memory CD4^+^ CD44^high^ T cells from the same condition (Supplementary figure 4). This PD-1 up-regulation, rather the Foxp3 expression, could reflect a high antigen reactivity and suggest the emergence of pro-inflammatory CD4 T cells.

Plasma levels of aminotransferases (AST/ALT) were used to monitor liver damage. In tamoxifen-fed mice and Ad Ct/Ad Cre i.m. groups, AST/ALT levels remained stable for 14 days and histological analysis of the liver did not show any sign of lymphocytic infiltration (Figure 1F). When adenoviral vectors (Cre or Ct) are injected intravenously, plasma AST/ALT levels were increased, with a peak after 7 days reflecting liver inflammation due to viral infection (Figure 1F). Interestingly, the histological analysis of the liver showed that portal infiltrations and liver damages were more severe in the liver of Ad Cre i.v. mice than in Ad Ct i.v. mice. Liver of Ad Cre i.v. mice showed many events of hepatocyte necrosis and a necroinflammatory activity (Figure 1F). Therefore, although the adenoviral vector itself induces a hepatic inflammation, the presence of HA-specific CD4^+^ CD4^high^ T cells associated with HA expression in the liver of Ad Cre i.v. mice could explain the exacerbated liver damages compared to Ad Ct i.v. mice.

Thus, the induction of the hepatic expression of the antigen HA in association with a concomitant adenoviral-mediated hepatic inflammation leads to the development of an antigen-specific response, marked by the generation of anti-HA antibodies and HA-specific CD4^+^ CD44^high^ PD-1^+^ T cells in the spleen and the liver. The presence of HA-specific CD4+ CD44^high^ PD-1+ T cells in the liver is associated with increased hepatic inflammation and liver damages, which could be explained by an immune attack against the HA-expressing hepatocytes, mimicking the early events of an autoreactive response.

### Antigen expression in the liver post-peripheral immunization leads to intra-hepatic recruitment of antigen-specific CD4^+^ T cells

HA-specific CD4^+^ T cells are observed in the liver of mice only after Ad Cre i.v. injection, but, it was not clear if these cells are generated within the liver or recruited from the periphery. We hypothesized that peripherally activated HA-specific memory T cells could be recruited to the liver following local HA expression and mediate local damages. For this, we analysed the consequences of a tamoxifen-induced hepatic expression of the antigen HA in peripherally pre-immunized mice. HA^fl^/Cre^ind-^ and HA^fl^/Cre^ind+^ mice were immunized by a single i.m. injection of Ad Cre. After 14 days, mice were fed with tamoxifen dry food for 14 days (Figure 2A).

**Figure 2:**
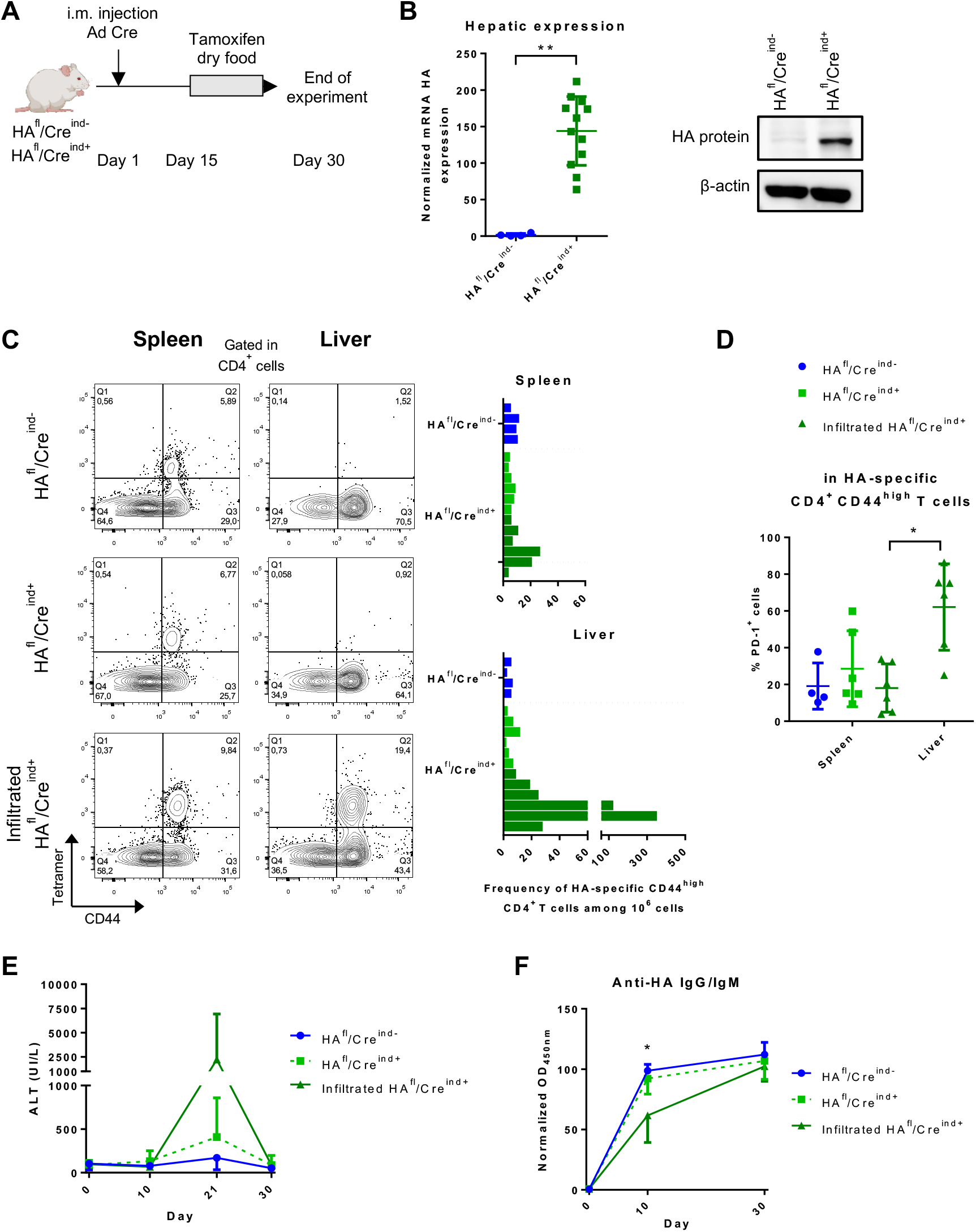
Induction of antigen expression in the liver of pre-immunized mice leads to intra-hepatic recruitment of antigen-specific CD4^+^ T cells. (A) HA^fl^/Cre^ind-^ mice (n = 4) and HA^fl^/Cre^ind+^ mice (n = 12) receive a single i.m. injection of Ad Cre (1,5.10^9^ ip). From day 15 to day 30, mice are fed with tamoxifen dry food (0,5g/kg). Mice are sacrificed at day 30. (B) Quantitative RT-PCR analysis of HA mRNA expression in liver samples. Total protein from liver samples were incubated on western blot membranes with anti-HA antibodies to detect HA protein expression. β-actine was used as loading control. (C) Total splenocytes and liver NPCs are stained with a MHC class II tetramer loaded with HA peptide before a tetramer enrichment step. Cells are stained (Live/Dead Aqua, CD4, CD44, PD-1, CD25, CD19) and analysed by flow cytometry. Representative MHC class II/HA peptide tetramer and CD44 co-staining, gated in live CD19^-^ CD4^+^ cells. Frequency of HA-specific CD4^+^ CD44^high^ T cells are calculated among 10^6^ total cells (grey and black bars respectively represent HA^fl^/Cre^ind+^ mice without or with antigen-specific CD4^+^ CD44^high^ T cells infiltration in the liver). (D) Percentage of PD-1^+^ cells in HA-specific CD4^+^ CD44^high^ T cells are represented. (E) Plasma samples were collected before the start of the experiment and at days 10, 21 and 30 for dosage of ALT. (F) Before the start of experiment, at day 10 and at the end of experiment, sera were diluted and assayed for anti-HA IgG and IgM. All results are normalized and represent the mean (+ SD). p values were calculated using non-parametric Kruskal-Wallis test or one-way ANOVA, ns = no significance, * = p <0.05, ** = p <0.01.

As expected, tamoxifen induced HA expression only in the liver of HA^fl^/Cre^ind+^ mice (Figure 2B). HA-specific CD4^+^ CD44^high^ T cells were detected in spleen of all groups of mice at similar frequencies (per 10^6^ cells: 8.75 ± 2.63 in HA^fl^/Cre^ind-^ mice; 8.50 ± 7.18 in HA^fl^/Cre^ind+^ mice), which confirms that all mice were indeed pre-immunized (Figure 2C). Tamoxifen induction of HA hepatic expression only in HA^fl^/Cre^ind+^ mice leads to a massive hepatic infiltration of HA-specific CD4^+^ CD44^high^ T cells in 50% of these mice (per 10^6^ cells: 89.67 ± 131.10 in liver-infiltrated HA^fl^/Cre^ind+^ mice, vs 4.83 ± 3.66 in non-infiltrated HA^fl^/Cre^ind+^ mice and 4.50 ± 1.73 in HA^fl^/Cre^ind-^ mice) (Figure 2C). In the infiltrated liver of HA^fl^/Cre^ind+^ mice, HA-specific CD4^+^ CD44^high^ PD-1^+^ T cells were enriched compared to their splenic counterpart (spleen: 18.06 ± 13.11%; liver: 62.13 ± 23.49%) (Figure 2D), while frequency of splenic and hepatic total memory CD4^+^ CD44^high^ PD-1^+^ T cells were still low (Supplementary figure 5). A hepatitis was associated with this antigen-specific immune response in liver-infiltrated HA^fl^/Cre^ind+^ mice, as shown by the increase in AST/ALT levels after HA hepatic expression induction (Figure 2E) and a residual diffuse infiltrate in the liver at the end of experiment (Supplementary figure 6). The humoral response, as measured by serum levels of anti HA IgG/IgM was comparable in mice of all groups at the end of the experiment. However, it is noteworthy that the mice that developed hepatitis associated with presence of liver HA-specific CD4^+^ T cells after tamoxifen diet are those in which the pre-immunisation had led to the weakest humoral immune response (HA^fl^/Cre^ind+^ mice: 92.25 ± 12.87 a.u. in non-infiltrated; 61.60 ± 22.32 a.u. in liver-infiltrated) (Figure 2F).

Thus, the expression of an antigen in the liver under pre-immunization condition is a factor to induce a local HA-specific CD4^+^ T cells recruitment associated to hepatitis, explaining the presence of HA-specific CD4^+^ T cells in the liver after i.v. Ad Cre infection as shown in Figure 1. These data also suggest that a weak pre-immunisation may be necessary to induce this local recruitment.

### Hepatic antigen expression induced by an adenovirus leads to a long-term peripheral antigen-specific response but local liver tolerance

In the first part of this work, we studied the first event involved in the generation and recruitment of HA-specific CD4^+^ T cell in the liver. However, liver autoimmunity is a long-term chronic disease in human. Thus, HA^fl^/Cre^ind-/+^ mice received either a unique i.v. injection of Ad Cre or of Ad Ct and were monitored for 24 weeks (Figure 3A).

**Figure 3:**
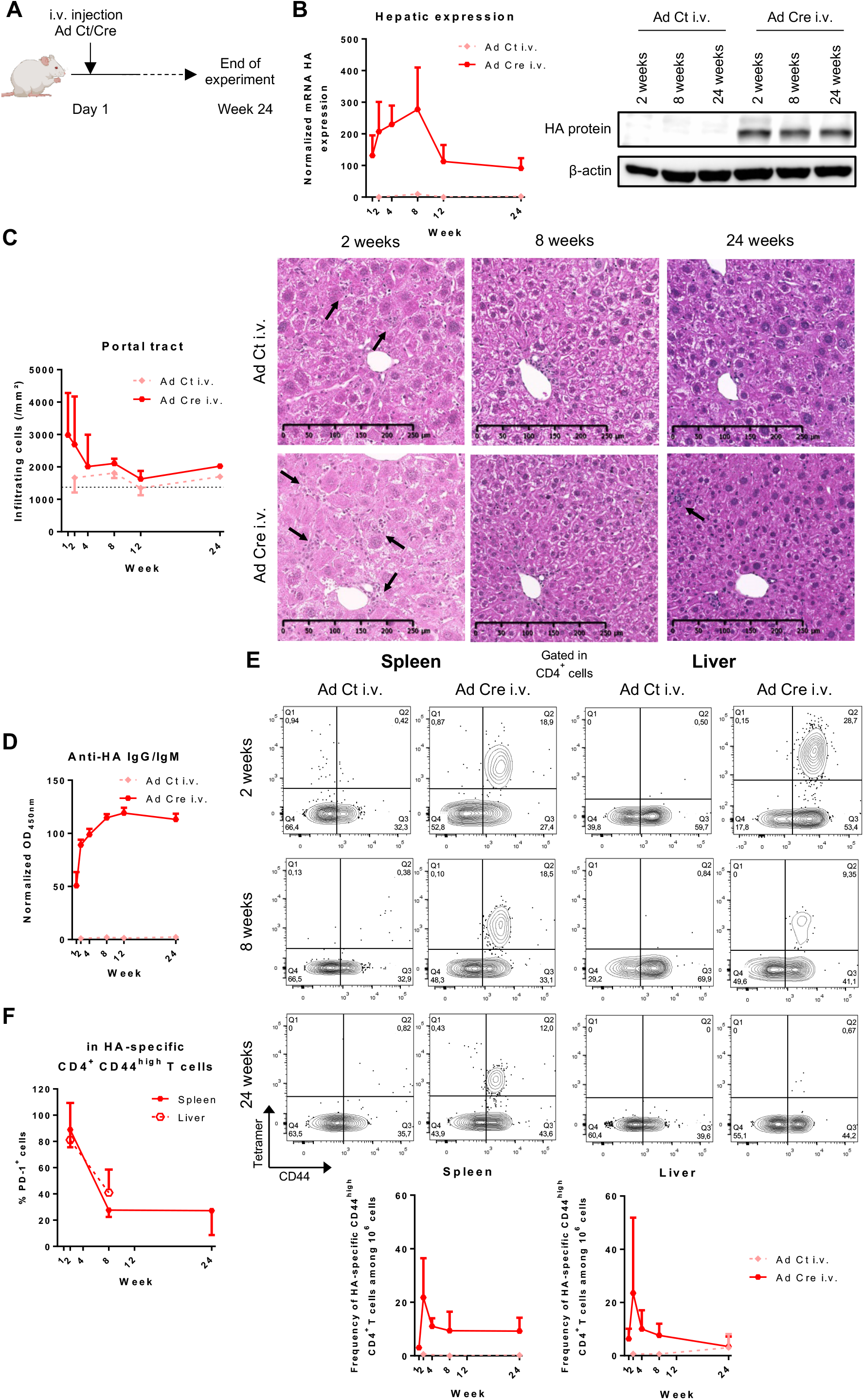
Induction of an antigen expression in the liver with concomitant inflammation using an adenoviral vector induces a chronic antigen-specific response. (A) HA^fl^/Cre^ind-/+^ mice receive a single i.v. injection (3.10^9^ ip) of Ad Ct or Ad Cre. Mice are sacrificed at week 1 (Ad Cre n = 3), 2 (Ad Ct n = 7; Ad Cre n = 13), 4 (Ad Cre n = 3), 8 (Ad Ct n = 8; Ad Cre n = 12), 12 (Ad Ct n = 3; Ad Cre n = 6) and 24 (Ad Ct n = 5; Ad Cre n = 9). (B) Quantitative RT-PCR analysis of HA mRNA expression in liver samples. Total protein from liver samples were incubated on western blot membranes with anti-HA antibodies to detect HA protein expression. β-actine was used as loading control. (C) For each time point, paraffin-embedded liver sections are stained with HPS coloration to analyse liver infiltration (count of infiltrating cells around portal tracts; dotted line represent a basal infiltration in a naïve HA^fl^/Cre^ind-/+^ mice) and morphology (x10; arrows point lymphocytic infiltration). (D) For each time point, sera were diluted and assayed for anti-HA IgG and IgM. (E) Total splenocytes and liver NPCs are stained with a MHC class II tetramer loaded with HA peptide before a tetramer enrichment step. Cells are stained (Live/Dead Aqua, CD4, CD44, PD-1, CD25 CD19) and analysed by flow cytometry. Representative MHC class II/HA peptide tetramer and CD44 co-staining, gated in live CD19^-^ CD4^+^ cells. Frequency of HA-specific CD4^+^ CD44^high^ T cells are calculated among 10^6^ total cells. (F) Percentage of PD-1^+^ cells in HA-specific CD4^+^ CD44^high^ T cells are represented. All results are normalized and represent the mean (+ SD)

24 weeks after i.v. injection of Ad Cre, expression of HA was still detectable in the liver (Figure 3B). Rapidly the hepatic damages are controlled in our mouse models. The peak of AST/ALT transaminases plasma levels observed at 1 week decreased to return to a basal level after 8 weeks (Ad Ct i.v.: 124.96 ± 71.69 UI/L; Ad Cre i.v.: 133,23 ± 83,66 UI/L) (data not shown). Likewise, liver infiltration and damages in Ad Cre i.v. mice decreased and were resolved after 4 weeks (Figure 3C). This is also associated with a decrease of HA-specific CD4^+^ T cells in the liver which suggest a local clearance over time (Figure 3E).

Interestingly, the global peripheral HA-specific adaptive immune response was only mildly impacted by the persistent HA expression in the liver. Anti-HA IgG and IgM were still detectable 24 weeks after injection, with a plateau reached after 8 weeks (Figure 3D). HA-specific CD4^+^ CD44^high^ T cells in Ad Cre i.v. mice were still detectable in the spleen at 24 weeks (Figure 3E). Long-lasting HA-specific CD4^+^ CD44^high^ T cells were still CD25- PD-1^+^, although frequency of PD-1^+^ cells decreased 2 weeks after Ad Cre injection (Figure 3F, Supplementary figure 7, Supplementary figure 8).

These data demonstrate a local HA-tolerance over time, but a limited impact on the peripheral pool of antigen-specific CD4^+^ T cells. The persistence of a peripheral pool of reactive CD4^+^ T cells could be involved in the chronic inflammatory events with the implication of other additional immune events.

## Discussion

In this study, we report that the induction of the expression of a model antigen by hepatocytes in non-inflammatory condition leads to antigen tolerance. The addition of an inflammatory cue, with an adenoviral vector, is sufficient to overcome immune tolerance and provoke hepatic local antigen reactivity. This leads to the generation of antigen-specific CD4^+^ T cells in the spleen and in the liver and is associated with mild liver damages which mimics a possible first step of local autoreactivity.

The different murine models of type 2 AIH directly target autoantigen expression in the liver using i.v. injection of an adenoviral vector encoding human FTCD or human CYP2D6. This leads to induction of an autoreactive response characterized by an antibody production and an hepatitis mediated by Th1 and Th17 cells^20,21^. While these models directly target the autoantigen via a mechanism of molecular mimicry, they lack in depth characterization of the antigen-specific T cell responses. Here, using an indirect antigen induction method (Ad Cre leading to HA expression), first we demonstrate that the expression of a neo-antigen in the liver in an immune-enhanced environment caused by an adenoviral vector is sufficient to generate an antigen-specific response, and second this response is marked by the generation of specific antibodies and antigen-specific CD4^+^ T cells in the spleen and in the liver. As for FTCD and CYP2D6 murine models, the presence of antigen-specific CD4^+^ T cells in the antigen-expressing liver in our model is associated with hepatic inflammation and hepatic damages, suggesting the initiation of an autoreactive response. This confirms that our murine models are accurate tools to further study the early events implicated in the generation of autoreactive CD4^+^ T cells in AIH.

Using two different models of induction of hepatic antigen expression, we show that antigen expression in the liver, either concomitant to a local inflammation cue (Ad Cre i.v.) or following a peripheral immunization (Ad Cre i.m. + Tamoxifen), mediates the hepatic recruitment of antigen-specific CD4^+^ T cells. This raises the question of the first site of the immune reactivity against the antigen in AIH, either within the liver or peripherally. The previous studies using AIH mouse models did not allow to conclude on this point. In our model, we observed a weak HA expression in the spleen after Ad Cre i.v. injection which could be the source of a peripheral immunisation. Thus, a concomitant antigen expression in the liver with the hepatic inflammation could explain the local accumulation and expansion of CD4^+^ T cells after peripheral generation.

Despite autoreactive response development, the HA-specific CD4^+^ T cell infiltrate in the liver disappears over time concurrently to hepatitis resolution, demonstrating a control of the autoreactive CD4^+^ T cell population in the liver allowing a long-term tolerance of antigen expression. However, the humoral response and frequency of peripheral antigen-specific CD4^+^ T cells are not affected in our model, despite the persistent expression of the HA antigen in the liver. These data suggest the HA specific immune response is only controlled locally and that additional immune events may be required within the liver to trigger a chronic hepatitis in this model.

To conclude, autoreactive CD4^+^ T cells are central to the immunopathogenesis of AIH. These are recruited in the liver following hepatic antigen exposure and mediate hepatic damages. In mice, a local control of this cellular autoreactive response rapidly emerges allowing a long-term tolerance of antigen expression, without chronic hepatitis. However, a pool of autoreactive CD4^+^ T cell persists in the spleen and could represent an active precursor of a chronic hepatitis. This murine model can be presented as an interesting pre-clinical model for the analysis of immunomodulatory molecules and pathways implicated in the local control of autoreactive CD4^+^ T cells. The identification of potential extrinsic factors implicated in an acute-to-chronic transition could provide a better understanding of AIH initiation mechanisms and unveil new therapeutical targets to dampen immunopathogenesis of autoreactive CD4^+^ T cells in the liver of patients.

## Abbreviations

AIH: autoimmune hepatitis
ALT: alanine transaminase
AST: aspartate transaminase
CYP2D6: cytochrome P450 2D6
FTCD: Formiminotransferase cyclodeaminase
HA: hemagglutinin
IgG: immunoglobulin G
i.m: intramuscular
i.v: intravenous
LC-1: liver cytosol 1
LKM-1: liver kidney microsomal type 1
MHC(-II): (class II) major histocompatibility complex
NPCs: non-parenchymal cells
SLA: soluble liver antigen
Tregs: regulatory T cells
TTR: transthyretin

## Acknowledgments

We are grateful to Pr. Liblau and Dr M Vasseur-Cognet for sharing the Rosa26 HA floxed and the TTR-Cre inducible mice strains.

We thank the NIH Tetramer Core Facility (contract number 75N93020D00005) for providing the class II HA-specific tetramers.

We wish to thank the UTE IRS-UN animal facility of the SFR Santé François Bonamy (Nantes Université, INSERM UAR016, CNRS UAR3556).

We acknowledge the IBISA MicroPICell facility (Biogenouest), member of the national infrastructure France-Bioimaging supported by the French national research agency (ANR-10- INBS-04).

## References

1. Kubes, P. & Jenne, C. Immune Responses in the Liver. Annu. Rev. Immunol. 36, 247–277 (2018).

2. Bénéchet, A. P. et al. Dynamics and genomic landscape of CD8+ T cells undergoing hepatic priming. Nature 574, 200–205 (2019).

3. Wherry, E. J., Blattman, J. N., Murali-Krishna, K., van der Most, R. & Ahmed, R. Viral Persistence Alters CD8 T-Cell Immunodominance and Tissue Distribution and Results in Distinct Stages of Functional Impairment. J. Virol. 77, 4911–4927 (2003).

4. Carambia, A. et al. TGF-β-dependent induction of CD4+CD25+Foxp3+Tregs by liver sinusoidal endothelial cells. J. Hepatol. 61, 594–599 (2014).

5. Heymann, F. et al. Liver inflammation abrogates immunological tolerance induced by Kupffer cells. Hepatol. Baltim. Md 62, 279–291 (2015).

6. Holz, L. E. et al. Intrahepatic murine CD8 T-cell activation associates with a distinct phenotype leading to Bim-dependent death. Gastroenterology 135, 989–997 (2008).

7. Benseler, V. et al. Hepatocyte entry leads to degradation of autoreactive CD8 T cells. Proc. Natl. Acad. Sci. 108, 16735–16740 (2011).

8. Cardon, A., Conchon, S. & Renand, A. Mechanisms of autoimmune hepatitis. Curr. Opin. Gastroenterol. 37, 79–85 (2021).

9. Sahebjam, F. & Vierling, J. M. Autoimmune hepatitis. Front. Med. 9, 187–219 (2015).

10. Manns, M. P. et al. Diagnosis and management of autoimmune hepatitis. Hepatology 51, 2193–2213 (2010).

11. Francque, S., Vonghia, L., Ramon, A. & Michielsen, P. Epidemiology and treatment of autoimmune hepatitis. Hepatic Med. Evid. Res. 4, 1–10 (2012).

12. Renand, A. et al. Immune Alterations in Patients With Type 1 Autoimmune Hepatitis Persist Upon Standard Immunosuppressive Treatment. Hepatol. Commun. 2, 968–981 (2018).

13. Behfarjam, F., Nasseri-Moghaddam, S. & Jadali, Z. Enhanced Th17 Responses in Patients with Autoimmune Hepatitis. Middle East J. Dig. Dis. 11, 98–103 (2019).

14. Bovensiepen, C. S. et al. TNF-Producing Th1 Cells Are Selectively Expanded in Liver Infiltrates of Patients with Autoimmune Hepatitis. J. Immunol. Baltim. Md 1950 203, 3148–3156 (2019).

15. Liang, M., Liwen, Z., Yun, Z., Yanbo, D. & Jianping, C. The Imbalance between Foxp3+Tregs and Th1/Th17/Th22 Cells in Patients with Newly Diagnosed Autoimmune Hepatitis. Journal of Immunology Research vol. 2018 e3753081 https://www.hindawi.com/journals/jir/2018/3753081/ (2018).

16. Longhi, M. S. et al. Impairment of CD4+CD25+ regulatory T-cells in autoimmune liver disease. J. Hepatol. 41, 31–37 (2004).

17. Oo, Y. H. et al. Liver homing of clinical grade Tregs after therapeutic infusion in patients with autoimmune hepatitis. JHEP Rep. Innov. Hepatol. 1, 286–296 (2019).

18. Peiseler, M. et al. FOXP3+ regulatory T cells in autoimmune hepatitis are fully functional and not reduced in frequency. J. Hepatol. 57, 125–132 (2012).

19. Renand, A. et al. Integrative molecular profiling of autoreactive CD4 T cells in autoimmune hepatitis. J. Hepatol. 73, 1379–1390 (2020).

20. Hardtke-Wolenski, M. et al. Genetic predisposition and environmental danger signals initiate chronic autoimmune hepatitis driven by CD4+ T cells. Hepatol. Baltim. Md 58, 718–728 (2013).

21. Holdener, M. et al. Breaking tolerance to the natural human liver autoantigen cytochrome P450 2D6 by virus infection. J. Exp. Med. 205, 1409–1422 (2008).

22. Tannour-Louet, M., Porteu, A., Vaulont, S., Kahn, A. & Vasseur-Cognet, M. A tamoxifen-inducible chimeric Cre recombinase specifically effective in the fetal and adult mouse liver. Hepatol. Baltim. Md 35, 1072–1081 (2002).

23. Saxena, A. et al. Cutting edge: Multiple sclerosis-like lesions induced by effector CD8 T cells recognizing a sequestered antigen on oligodendrocytes. J. Immunol. Baltim. Md 1950 181, 1617–1621 (2008).

24. Le Guen, V. et al. Alloantigen gene transfer to hepatocytes promotes tolerance to pancreatic islet graft by inducing CD8(+) regulatory T cells. J. Hepatol. 66, 765–777 (2017).

