## supplementary figures for "Preclinical model for the study of immune responses specific for a hepatic self-antigen"

† Equal contribution

### Supplementary figures

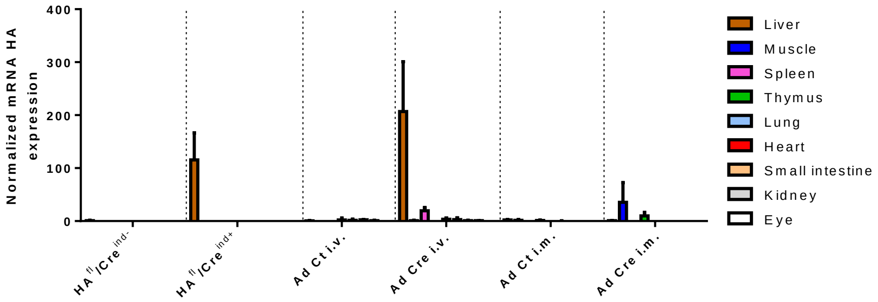

**Supplementary figure 1 : mRNA HA expression in different organs.** HA$^{fl}$/Cre$^{ind-}$ mice (n = 5) and HA$^{fl}$/Cre$^{ind+}$ mice (n = 9) are fed with tamoxifen dry food (0,5g/kg) for 14 days. HA$^{fl}$/Cre$^{ind-/+}$ mice receive a single i.v. injection ($3.10^9$ ip) of Ad Ct (n = 7) or Ad Cre (n = 9). HA$^{fl}$/Cre$^{ind-/+}$ mice receive a single i.m. injection ($1,5.10^9$ ip) of Ad Ct (n = 3) or Ad Cre (n = 9). Mice are sacrificed at day 15. Quantitative RT-PCR analysis of mRNA HA expression in different organs. All results are normalized and represent the mean (+ SD).

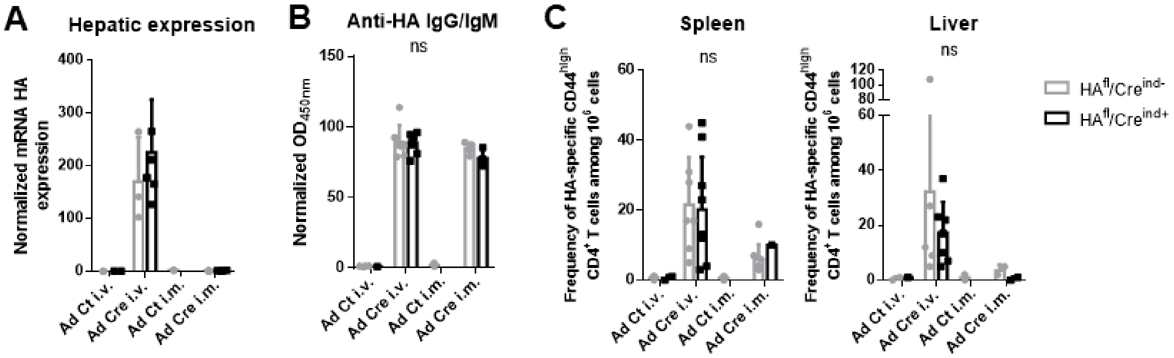

**Supplementary figure 2 : Effects of Ad Cre (i.v. or i.m.) in HA$^{fl}$/Cre$^{ind-}$ mice and HA$^{fl}$/Cre$^{ind+}$ mice are similar.** HA$^{fl}$/Cre$^{ind-}$ mice and HA$^{fl}$/Cre$^{ind+}$ mice are used indiscriminately for Ad Cre protocols (i.v. or i.m.) and show similar levels of (A) mRNA HA levels, (B) anti-HA IgG and IgM and (C) HA-specific CD4$^{+}$ CD44$^{high}$ T cells in spleen and liver. All results represent the mean (+ SD). p values were calculated using non-parametric Kruskal-Wallis test between HA$^{fl}$/Cre$^{ind-}$ mice and HA$^{fl}$/Cre$^{ind+}$ mice in each condition, ns = no significance.

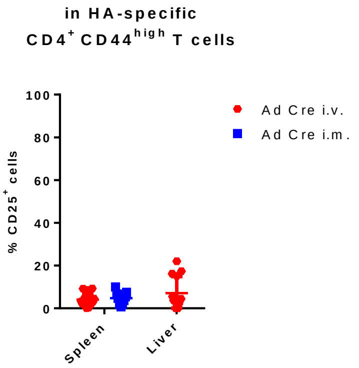

**Supplementary figure 3 : Phenotyping of HA-specific CD4$^+$ CD44$^{high}$ T cells induced by an i.v. or an i.m. Ad Cre injection.** Total splenocytes and liver NPCs are stained with a MHC class II tetramer loaded with HA peptide before a tetramer enrichment step. Cells are stained (Live/Dead Aqua, CD4, CD44, PD-1, CD25, FoxP3, CD19) and analysed by flow cytometry. Percentage of CD25$^+$ cells in HA-specific CD4$^+$ CD44$^{high}$ T cells are represented. Results represent the mean (+ SD).

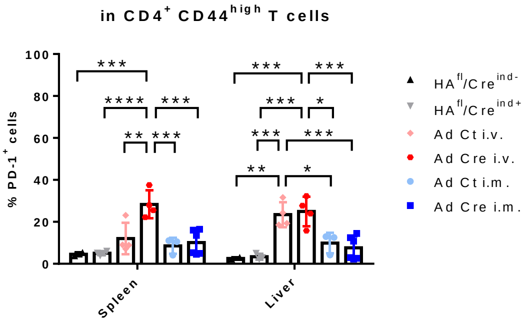

**Supplementary figure 4 : PD-1 expression in CD4$^+$ CD44$^{high}$ T cells.** HA$^{fl}$/Cre$^{ind-}$ mice (n = 2) and HA$^{fl}$/Cre$^{ind+}$ mice (n = 4) are fed with tamoxifen dry food (0,5g/kg) for 14 days. HA$^{fl}$/Cre$^{ind-/+}$ mice receive a single i.v. injection (3.10$^9$ ip) of Ad Ct (n = 4) or Ad Cre (n = 4). HA$^{fl}$/Cre$^{ind-/+}$ mice receive a single i.m. injection (1,5.10$^9$ ip) of Ad Ct (n = 3) or Ad Cre (n = 6). Mice are sacrificed at day 15. Total splenocytes and liver NPCs are stained with Live/Dead Aqua, CD4, CD44, PD-1, CD19 and analysed by flow cytometry. Percentage of PD-1$^+$ cells in spleen and liver are calculated among CD4$^+$ CD44$^{high}$ T cells populations. All results represent the mean (+ SD). p values were calculated using Tukey's multiple comparisons test, * = $p < 0.05$, ** = $p < 0.01$, *** = $p < 0.001$, **** = $p < 0.0001$.

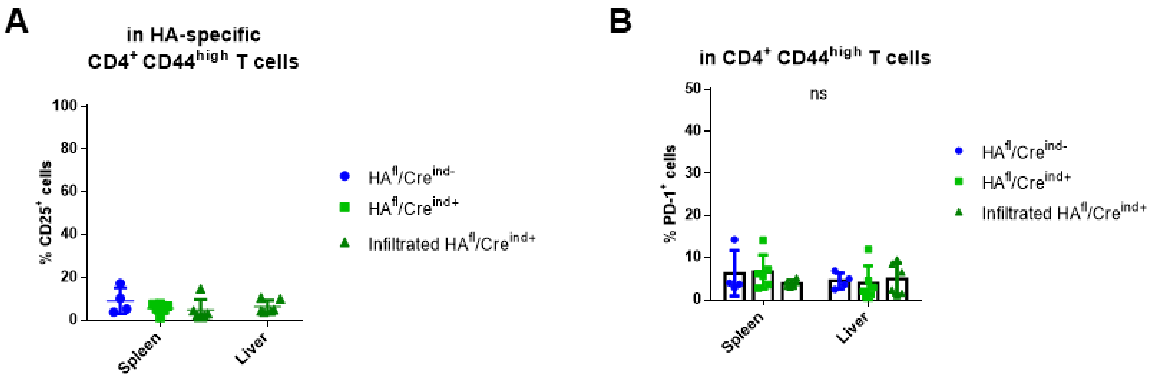

**Supplementary figure 5 : Phenotype of antigen-specific CD4$^+$ CD4$^{high}$ T cells and total memory CD4$^+$ CD44$^{high}$ T cells in tamoxifen-fed HA$^{fl}$/Cre$^{ind-}$ and HA$^{fl}$/Cre$^{ind+}$ pre-immunized mice.** HA$^{fl}$/Cre$^{ind-}$ mice (n = 4) and HA$^{fl}$/Cre$^{ind+}$ mice (n = 12) receive a single i.m. injection of Ad Cre (1,5.10$^9$ ip). From day 15 to day 30, mice are fed with tamoxifen dry food (0,5g/kg). Mice are sacrificed at day 30. Total splenocytes and liver NPCs are stained with Live/Dead Aqua, CD4, CD44, PD-1, CD25, CD19 and analysed by flow cytometry. (A) Percentage of CD25$^+$ cells in HA-specific CD4$^+$ CD44$^{high}$ T cells in spleen and liver are represented. (B) Percentage of PD-1$^+$ cells in spleen and liver are calculated among CD4$^+$ CD44$^{high}$ T cells population. All results represent the mean (+ SD). p values were calculated using Tukey's multiple comparisons test, ns = no significance.

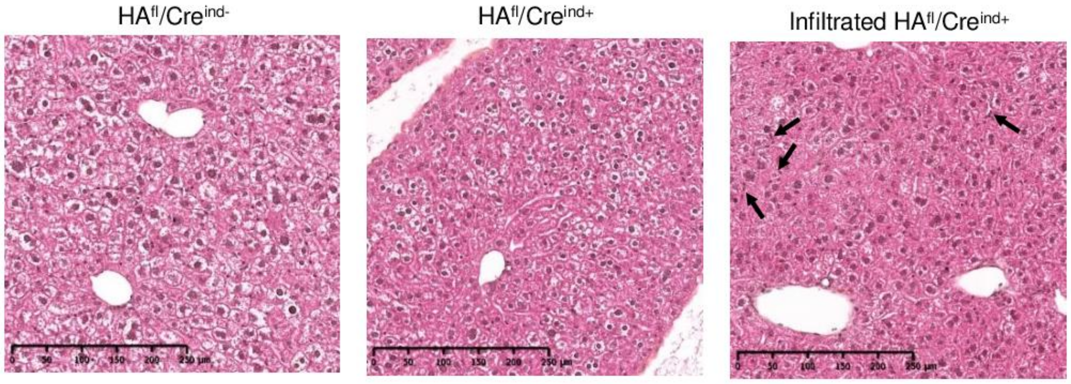

**Supplementary figure 6 : Liver-infiltrated HA$^{fl}$/Cre$^{ind+}$ mice show a residual diffuse infiltrate.** At the end of the experiment, paraffin-embedded liver sections are stained with HPS coloration to analyse liver morphology (x10 ; arrows point lymphocytic infiltration).

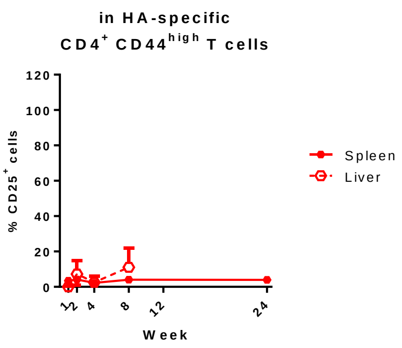

**Supplementary figure 7 : Phenotyping of HA-specific CD4$^+$ CD44$^{high}$ T cells induced by an i.v. Ad Cre injection.** Total splenocytes and liver NPCs are stained with a MHC class II tetramer loaded with HA peptide before a tetramer enrichment step. Cells are stained (Live/Dead Aqua, CD4, CD44, PD-1, CD25, CD19) and analysed by flow cytometry. Percentage of CD25$^+$ cells in HA-specific CD4$^+$ CD44$^{high}$ T cells are represented. Results represent the mean (+ SD).

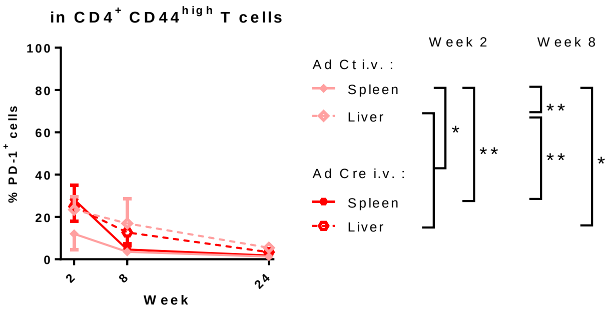

**Supplementary figure 8 : PD-1 expression in CD4$^+$ CD44$^{high}$ T cells after i.v. injection of Ad Ct or Ad Cre.** HA$^{fl}$/Cre$^{ind-/+}$ mice receive a single i.v. injection (3.10$^9$ ip) of Ad Ct or Ad Cre. Mice are sacrificed at 2 (Ad Ct n = 4 ; Ad Cre n = 4), 8 (Ad Ct n = 6 ; Ad Cre n = 6) and 24 (Ad Ct n = 4 ; Ad Cre n = 5) weeks. Total splenocytes and liver NPCs are stained with Live/Dead Aqua, CD4, CD44, PD-1, CD19 and analysed by flow cytometry. Percentage of PD-1$^+$ cells in spleen and liver are calculated among CD4$^+$ CD44$^{high}$ T cells population. All results represent the mean (+ SD). p values were calculated using Tukey's multiple comparisons test, * = p <0.05, ** = p <0.01.
